# Focused ultrasound-mediated brain genome editing

**DOI:** 10.1101/2022.12.10.519924

**Authors:** Yeh-Hsing Lao, Robin Ji, Joyce K. Zhou, Kathy J. Snow, Nancy Kwon, Ethan Saville, Siyu He, Shradha Chauhan, Chun-Wei Chi, Malika S. Datta, Hairong Zhang, Chai Hoon Quek, Sarah Cai, Mingqiang Li, Yaned Gaitan, Lawrence Bechtel, Shih-Ying Wu, Cathleen M. Lutz, Raju Tomer, Stephen A. Murray, Alejandro Chavez, Elisa E. Konofagou, Kam W. Leong

## Abstract

Gene editing in the mammalian brain has been challenging because of the restricted transport imposed by the blood-brain barrier (BBB). Current approaches rely on local injection to bypass the BBB. However, such administration is highly invasive and not amenable to treating certain delicate regions of the brain. We demonstrate a safe and effective gene editing technique by using focused ultrasound (FUS) to transiently open the BBB for the transport of intravenously delivered CRISPR/Cas9 machinery to the brain.

---

CRISPR gene editing technologies provide exciting opportunities to advance gene therapy and treat many intractable genetic diseases, including neurodegenerative disorders.^1,2^ However, effective *in vivo* delivery of CRISPR components remains a significant barrier. Except for the liver, CRISPR delivery mainly relies on local administration. Given its highly invasive nature, this can be particularly problematic when targeting the brain. A safer yet effective method of delivery would help empower the use of somatic gene editing in this critical organ. We and other groups have demonstrated FUS delivery of drugs and biologics to the brain through systemic routes.^3^ Our efforts have led to a clinical trial (NCT04804709) for delivering Panobinostat to children with diffuse midline glioma in the brainstem, a difficult and sensitive region for direct administration. Here we report the feasibility of applying FUS to achieve gene editing in targeted brain regions following the intravenous injection of AAV vectors encoding CRISPR/Cas9 machinery.

FUS-mediated BBB opening is accomplished through the cavitation of systemically administered microbubbles in the FUS focus, temporarily permeabilizing the BBB at the FUS-targeted site for the delivery of various payloads (Fig. 1a). Based on the therapeutic goal, the target region, dose and FUS parameters can be modified to maximize the transport of AAV into the brain. We previously developed two different FUS systems, spherical single-element FUS^3^ and FUS array,^4^ enabling transient opening of the BBB in a more confined or widespread region, respectively (Fig. 1b). We first started with the spherical system to test whether FUS could reproducibly improve the delivery of AAV/*S. aureus* Cas9 (SaCas9) vector into the mouse brain. In adult C57BL/6 mice, we were able to reproducibly tune the BBB permeability at the same brain region (two openings in the left hemisphere: 4mm left/3mm above and 5mm left/2mm above, relative to lambda). Although AAV9 was reported to have CNS-tropism when given intravenously,^5^ its brain deposition was still significantly lower than the level seen in other organs on our hands. In contrast, FUS enhanced the deposition of SaCas9-encoding AAV9 by ∼13 times at the target hemisphere, thus allowing targeting of the brain at levels similar to other organs apart from the liver (Fig. 1c). We observed a similar enhancement when directly comparing the FUS-targeted hemisphere with its contralateral side in the same animal, in line with FUS allowing one to precisely control the region of the brain that will receive the cargo of interest (Extended Data Fig. 1).

**Fig. 1.**
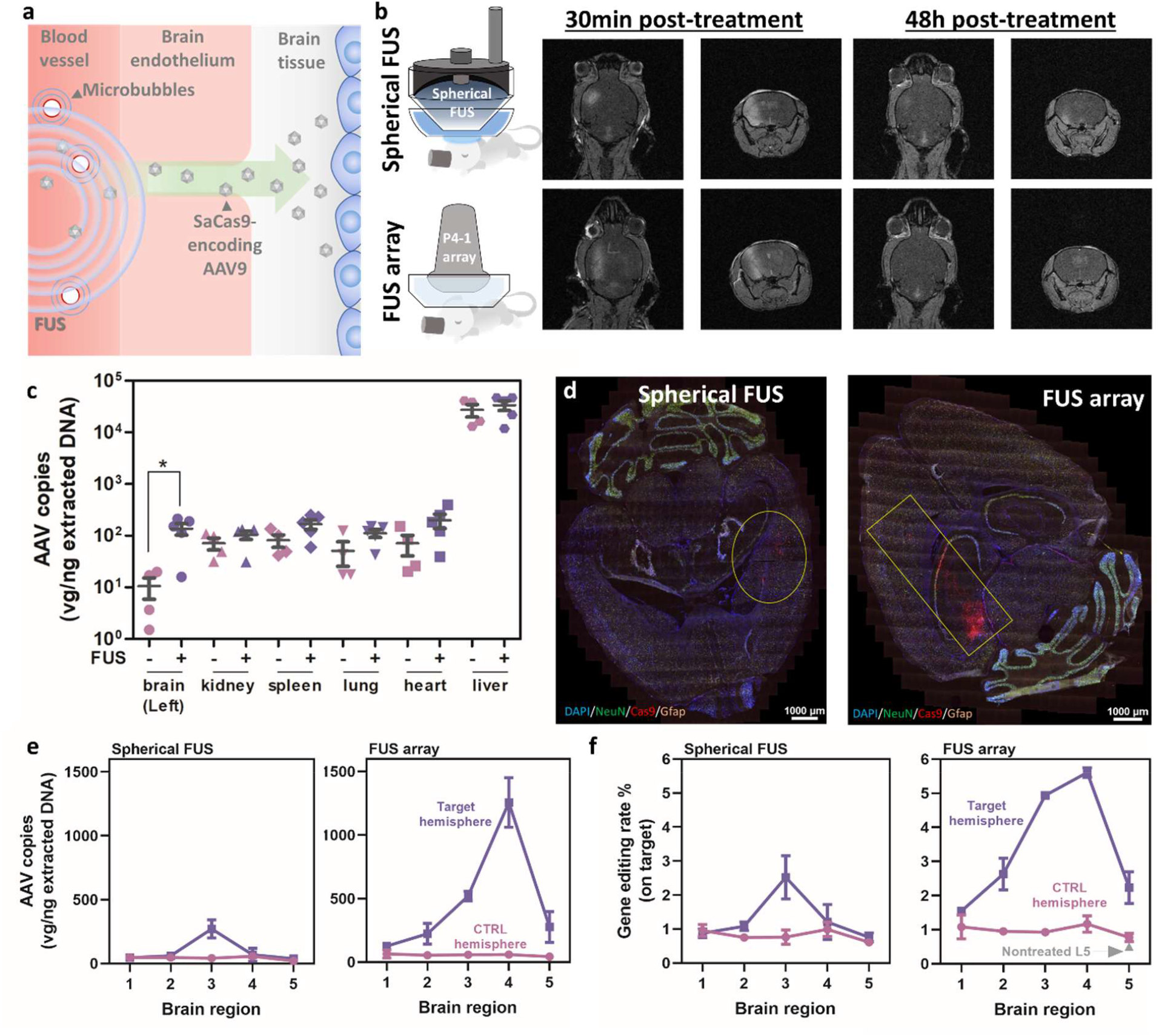
FUS to enhance systemic AAV/CRISPR vector delivery to the brain. **a**, Schematic overview of FUS-mediated BBB opening. **b**, Two types of FUS systems used in this study and the representative MRI images showing the transient opening induced by FUS. **c**, Biodistribution of SaCas9/AAV9 vector at week 2 post-administration. Both FUS and control groups received an AAV dose of 2×10^11^ GC/mouse (*N*=5 for the FUS group and *N*=4 for the control group, adult male C57BL/6). **d**, Representative RNAscope images to confirm the SaCas9 expression in the FUS-targeted region in adult C57BL/6 mice **e**, Deposition of SaCas9/AAV9 vector in different brain regions (Two biological repeats for each group). **f**, Gene editing efficiency in different brain regions (determined by amplicon sequencing; two biological repeats for each group). For Figs. 1d, e and f, adult C57BL/6 mice (aged between 9 to 10 weeks old) were given intravenously with SaCas9/AAV9 vectors in a dose of 10^12^ GC/mouse, and the brain tissues were dissected at week 3 post-administration.

We next optimized the SaCas9 vector by swapping the promoter and modifying the guide RNA (gRNA) scaffold as certain viral promoters (e.g., CMV) could be transcriptionally silenced in brain cells,^6^ while the poly-T motif in the wild-type gRNA scaffold may lead to gRNA early termination.^7,8^ As expected, the constitutive mammalian promoter EF1α enhanced *in vivo* SaCas9 expression by 11-fold when compared with its parental vector containing the CMV promoter (Extended Data Fig. 2a). Furthermore, the engineered variant of gRNA scaffold significantly improved gene editing efficiency *in vitro* after optimization (Extended Data Fig. 2b). We tested this optimized vector using a well-characterized *Pcsk9* guide^9^ to determine the biodistribution of our system when packaged into AAV and delivered systemically. At a dose of 2×10^11^ genome copies (GC)/mouse, we observed a significant reduction in total cholesterol levels (Extended Data Fig. 2c), consistent with the results reported in the literature.^9^ However, we only detected a minimal indel rate in the target locus from the brain samples (∼1%, Extended Data Fig. 2d).

**Fig. 2.**
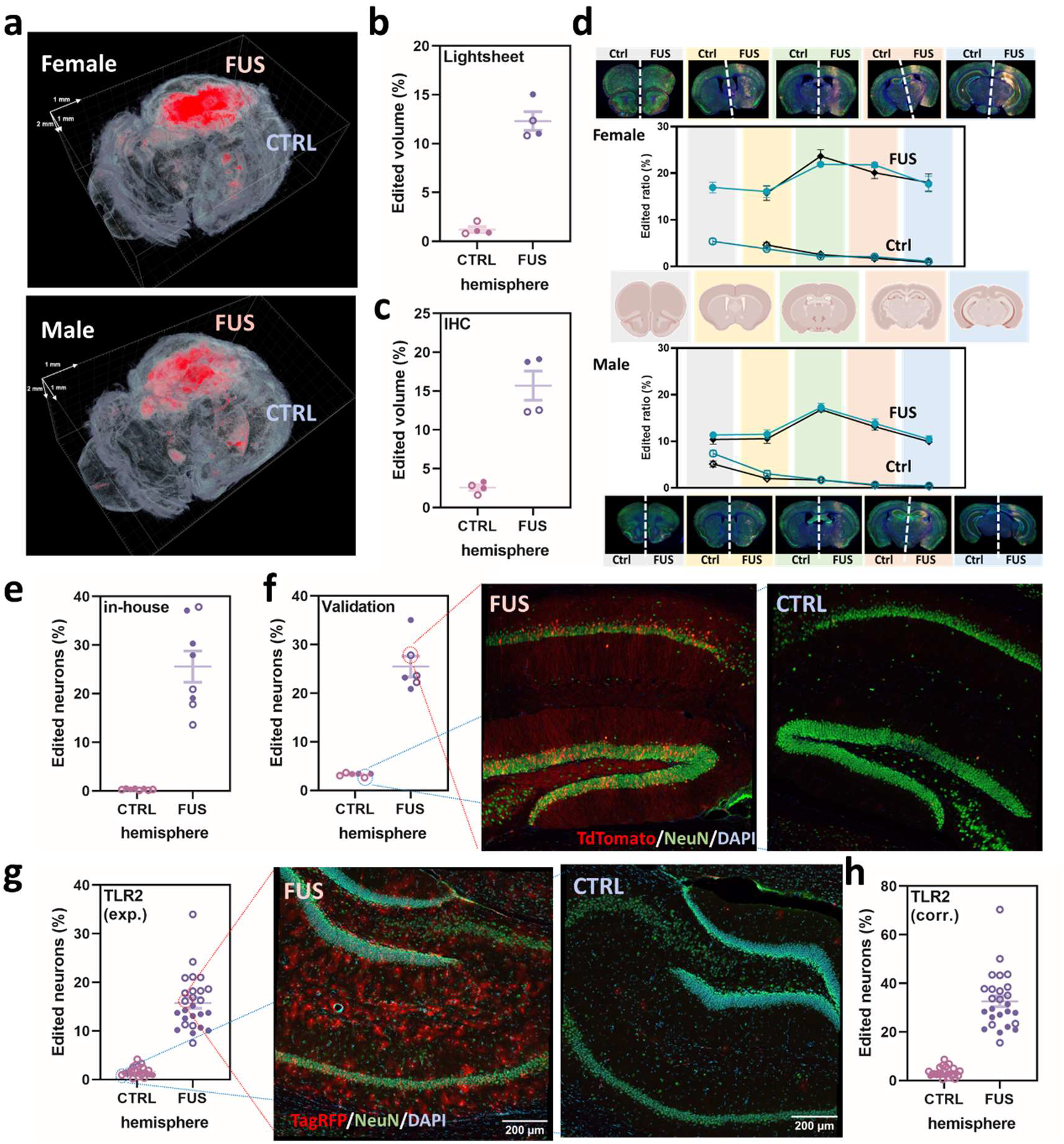
Validation of FUS-mediated brain gene editing in reporter mouse models. **a**, Representative reconstructed 3D Lightsheet images showing the Cas9-activated TdTomato signals (red) in the FUS-targeted regions. **b**, Gene editing efficiency quantified by a Lightsheet microscope. **c**, Efficiency quantification by immunostaining. For both Figs. 2b and c, Efficiency was determined by TdTomato+ volume per total volume of the hemisphere. **d**, Gene editing efficiency in different brain regions. Two sets of serial sections were used for immunostaining and quantification for each sex. **e**, In-house quantification of edited neurons (TdTomato+ and NeuN+) in the hippocampus in FUS-SaCas9/AAV9-treated Ai9 mice by immunostaining. **f**, Independent quantification of neuron editing in Ai9 mice following FUS performed by the SCGE Small Animal Testing Center and representative confocal images of the FUS-targeted and control sites. **g**, Quantification of TagRFP+ neurons in the hippocampus regions in FUS-SaCas9/AAV9-treated TLR2 reporter mice by immunostaining and the representative confocal images used for this quantification. **h**, Neuron editing performance determined with the experimental correction. Data are presented in dots and circles for the results from female and male mice, respectively.

Next, we increased the dose to 10^12^ GC/mouse and also included the FUS array system to see if gene editing in the brain could be significantly improved. We first established that this AAV dose was acceptable when administered intravenously, as blood chemistry analysis did not reveal any toxicity in C57BL/6 mice (Extended Data Fig. 3). At the FUS targeted regions in the brain, we could detect significant SaCas9 RNA transcripts from both FUS groups, and as aforementioned, the spherical FUS produced gene editing in a confined volume, while the FUS array led to Cas9 expression that was more widespread (Fig. 1d). For quantitative analysis, we further divided the brain tissues into five regions, and the qPCR and amplicon sequencing results matched the results from RNA hybridization. When comparing these two systems, we did see higher Cas9 vector deposition (>3×) in the FUS array group (Fig. 1e), leading to an enhanced gene editing efficiency (Fig. 1f). Overall, using the FUS array system with a systemic dose of 10^12^ GC/mouse, we were able to reach nearly 10% gene editing from this unbiased, bulk tissue analyses.

To further validate the gene editing efficacy, we tested our approach in the Ai9 mouse reporter line carrying a loxP-STOP-loxP-CAG-TdTomato cassette.^10^ Deletion of the loxP-flanked transcriptional terminator (STOP) by dual guide Cas9 editing can activate the downstream expression of TdTomato.^7^ Given the need of two gRNAs to completely remove the stop signal, an additional U6-gRNA expression cassette was added to our vector, and in the *in vitro* validation with an Ai9 reporter-containing HEK293T cell line, we confirmed that our dual-targeting vector could activate TdTomato (Extended Data Fig. 4a). In light of the use of high dose AAV (10^13^ GC) for Cas9 editing via intracranial administration setting in other work^11^ and potentially lower efficiency by requiring Cas9 to cut two targets, we chose 2×10^12^ GC/mouse as our systemic administration dose for these follow-up validations. Adult Ai9 mice received our Ai9-targeting vector intravenously under the FUS array system. At the endpoint (week 3 post-administration), we observed significant TdTomato activation at the target hemisphere when tissue was examined using a Lightsheet microscope (Fig. 2a). Normalizing the TdTomato+ volume to the whole hemisphere, the overall editing efficiencies were determined to be 12.3% and 1.21% for the FUS-targeted and contralateral hemispheres, respectively (Fig. 2b). In parallel, we serially sectioned the mouse brains and carried out histological analysis. Aligned with the Lightsheet results, we saw significant gene editing in the FUS-targeted hemisphere with an efficiency of 15.7% (Fig. 2c). We grouped the sections based on their locations and quantified the average gene editing efficiency for each region. The editing performance profile in Ai9 correlated with the trend we saw from the sequencing results in C57BL/6 mice (Figs. 1f and 2d), and two sets of serial sections gave consistent results (Fig. 2d). We noticed stronger editing in the female mice, and this could be attributed to the weight difference between sexes at the same age (average 18.6g for female vs. 26.7g for male), which led to a different dose per weight in the two groups. Nevertheless, the use of FUS significantly improved the brain gene editing efficacy by enhancing the brain deposition of the AAV vectors. We then analyzed the editing efficiency in neurons in the hippocampus, where the center of our FUS array was positioned. In the selected regions-of-interest (ROIs: 1,200×1,200 μm^2^; the focal size of our FUS probe), we found 25.6% of the neurons edited in this particular region, while <1% observed in the contralateral control sites (Fig. 2e and Extended Data Fig. 4b).

Through participation in the NIH Somatic Cell Genome Editing (SCGE) Program,^12^ we worked with the SCGE Small Animal Testing Center (SATC) at the Jackson Laboratory (JAX Lab) to verify the effectiveness and reproducibility of our technology. The same analysis pipeline performed at the SATC showed a consistent result, 25.5% of the neurons edited in the hippocampus region in the Ai9 model (Fig. 2f). The SATC validated our approach using an independent Traffic Light Reporter model (TLR2) generated for the SCGE program. This reporter strain carries a mutated Venus-P2A-TagRFP cassette, where a double-strand break (DSB) in the reporter followed by a nonhomologous end-joining DNA repair event can activate TagRFP, and if a donor is provided, a homology-directed repair event can restore Venus expression.^13^ Since this model has not been used with SaCas9, we first optimized the guide sequence *in vitro* (Extended Data Fig. 5a). In our *in vivo* TLR2 validation, 15.8% of the neurons were TagRFP-positive in the hippocampus ROIs at the FUS array targeted site (Fig. 2g). Because only indels in the +1/-2 frame could activate the TagRFP expression, we established a correction factor (2.07) based on the amplicon sequencing results obtained from whole FUS-targeted hemisphere (Extended Fig. 5b) to estimate the actual overall editing efficiency (32.8%, Fig. 2h). Better performance seen in TLR2 may be because only one DSB is needed for TagRFP activation, which could be more efficient versus the two DSBs and deletion of the STOP cassette required for activation of Ai9. Altogether, the results in two mouse models across two different laboratories confirmed the robustness of the FUS technology for brain gene editing.

In summary, we demonstrate that FUS is a reproducible CRISPR delivery approach for effective gene editing in specific brain regions through systemic administration of CRISPR-encoding vectors. By combining FUS with AAV-mediated gene delivery, we can achieve >25% editing efficiency of particular cell types, which is noteworthy as our approach is still dependent upon the tropism of the AAV capsid used, which does not infect all neuron types equally, even within a narrow region of the brain. Furthermore, the efficiency of the gRNAs and the ability of the promoters to drive robust expression of the CRISPR components also play key roles in driving editing rates. In future studies, the efficiency of this FUS-based approach is likely to be enhanced by further engineering the carrier and the CRISPR components. Our previous studies in larger animals^14-16^ and human trials (NCT04804709 and NCT04118764) have proven the safety and applicability of FUS. The method established here has the potential to expand the toolkit options for CRISPR delivery and open new opportunities for treating diseases of the brain, such as neurodegenerative disorders, with somatic genome editing.

## Online Methods

All animal experiments were conducted in compliance with the protocols AC-AABD2600 (animal breeding), AC-AABD5600 and AC-AABG4559 (FUS experiments), which were approved by the Institutional Animal Care and Use Committee at Columbia University.

### AAV vectors and production

Our SaCas9 AAV vectors were built based on the AAV2 ITR-flanked, CMV-driven SaCas9 AAV vector (Takara) by swapping the CMV promoter with the human EF1α promoter and the gRNA scaffold with the published variants.^7,8^ Our cloned vectors were verified by Sanger sequencing by Genewiz to confirm the whole sequences of SaCas9, U6-gRNA expression cassette and the ITR regions. To further clone the guide sequences, we followed the protocols from our previously published work.^17^ All the guides used in this study are listed in Supplementary Table 1. The plasmids will be deposited in Addgene and available to the community for other applications. The AAVs used in this study were either produced in-house (for the experiments done in C57BL/6 mice) or by PackGene (for the validations done in Ai9 and TLR2 mice). For in-house production, the SaCas9-encoding AAV vector, AAV2/9 Rep-Cap (obtained from the University of Pennsylvania Viral Vector Core Facility under the Materials Transfer Agreement) and the pHelper (Takara) plasmids were co-transfected to the HEK293T cell (Takara) using the CalFectin transfection reagent (SignaGen) in a ratio of 1:1:1. At 72h post-transfection, the cells were harvested, and the viruses were extracted using Takara AAV Purification Kit followed by buffer exchange using the AMICON-15 column (Millipore-Sigma, MWCO 100 kDa). After concentration, the virus was stored at -80°C in 1×PBS with 5% glycerol. Prior to use, the AAV concentration was quantified by qPCR using the quantification kit from Takara.

### *In vitro* validation

To determine which gRNA scaffold to use, three EF1a-SaCas9 AAV plasmids (containing wild-type or the optimized scaffold) targeting mouse *Pcsk9* (see Supplementary Table 1 for guide and scaffold sequences) were tested *in vitro*. Mouse Neuro-2a cell (ATCC) was first seeded in a 24-well plate (Corning-Falcon) with a density of 100,000 cells/well. After being cultured overnight, the AAV plasmid (500ng) was co-transfected with 100ng EGFP plasmid (Addgene# 46956) using Lipofectamine 3000 (Invitrogen) by following the manufacturer’s instructions. At 48h post-transfection, genomic DNAs were extracted using Zymo Quick-DNA MiniPrep Plus kit (Zymo Research), and the target Pcsk9 locus was amplified using Terra PCR Direct Polymerase (Takara; primers listed in Supplementary Table 2). After column purification, the concentration of the PCR product was quantified by PicoGreen (Thermo-Fisher), and amplicon NGS was done by Genewiz.

For the TLR2 guide screening, five top guides targeting the TLR2 locus were picked using the GPP sgRNA Designer^18^ (see Supplementary Table 1 for guide information) and tested *in vitro* in the TLR2-encoding HEK293T reporter cells (obtained from Max Delbrück Center for Molecular Medicine under the Materials Transfer Agreement). Similarly, the reporter cell was first seeded in a 24-well plate with a density of 150,000 cells/well, and the TLR2-targeting AAV plasmids were transfected using Lipofectamine 3000 on the second day. At 48h post-transfection, cells were harvested and TagRFP activation efficiency was quantified by flow cytometry. The genomic DNAs were then extracted followed by amplification at the TLR2 locus (primers listed in Supplementary Table 2) using the same aforementioned kit and polymerase. Amplicon NGS was performed by Genewiz, and the results were analyzed using CRISPresso2.

### FUS-mediated BBB opening

FUS-mediated BBB opening was achieved using two different types of transducer setups: (1) a spherical single-element transducer or (2) a phase array probe. First, mice were placed on a heating pad under anesthesia (1-2% isoflurane with oxygen at a flow rate of ∼0.8L/min) and fixed to a stereotactic frame (David Kopf Instruments). Afterwards, the head of the mouse was shaved, and depilatory cream was applied to remove any remaining hair. Degassed ultrasound gel was then applied to the mouse scalp to couple a chamber containing degassed and deionized water. For FUS-mediated BBB opening with a single-element transducer, a spherical FUS transducer (center frequency: 1.5 MHz; diameter: 60 mm, focal depth: 60 mm; Imasonic), controlled by the 3D positioning system (Velmex), was targeted to the caudate putamen in two spots. A sterile saline solution containing in-house made polydispersed microbubbles (8×10^8^ bubbles/mL)^19^ and AAVs were co-administered intravenously, followed by FUS sonication with a pulsed length of 1,000 cycles (0.67 ms) and a pulse repetition frequency (PRF) of 5 Hz at an estimated derated acoustic peak-negative pressure of 0.6 MPa. Immediately after the co-injection of microbubbles and AAVs, sonications were performed in the left striatum in two locations (relative to the cranial landmark lambda, target 1 - 4 mm anterior and 3 mm lateral, target 2 - 5 mm anterior, 2 mm lateral) consecutively for 1 minute each.

For BBB opening with the FUS array system, a P4-1 ultrasound phased array probe (ATL, Philips) was instead connected to the 3D positioning system and controlled by a Vantage system (Verasonics Inc.). To open the BBB, rapid bursts of short pulses (transmit frequency: 1.5 MHz, pulse length: ∼3-cycle pulses) were pulsed at a PRF of 1 kHz, with a total of 100 pulses transmitted per burst. The burst repetition frequency (BRF) was 0.5 Hz, and a total of 60 bursts were transmitted per treatment session, for a total treatment duration of 2 minutes. The FUS array was placed over the left hemisphere (relative to the cranial landmark lambda, 3 mm anterior and 2 mm lateral). The free-field peak negative pressure of the P4-1 focused transmits was estimated to be 1.5 MPa. Electronic delays were applied to set the transducer focus at a depth of 35 mm. Similarly, sonications were performed right after the intravenous administration of the saline solution containing microbubbles and AAVs.

Following each FUS procedure, the mouse was given 0.2 mL of gadodiamide (Omniscan, GE Healthcare) intraperitoneally, and the BBB opening was confirmed using a 9.4T MRI system (Bruker Biospin) through a contrast-enhanced T1-weighted 2D FLASH sequence (TR/TE: 230/3.3 ms; flip angle: 70°; number of excitations: 6; field of view: 25.6×25.6 mm^2^; resolution: 100×100×400 μm^3^).

### Gene editing analysis in C57BL/6 mice

Adult male C57BL/6 mice (9-10 weeks old; Envigo) were intravenously given AAV9 encoding *Pcsk9*-targeting SaCas9 with or without FUS treatment. At the endpoint (week 3 post-administration, if without specific mention), the mouse was transcardially perfused with cold 1×PBS under anesthesia. After harvesting, the whole mouse brain was directly embedding optimal cutting temperature (OCT) compound on a dry ice slurry for RNA hybridization, or two brain hemispheres were dissected into 5 regions using a mouse brain matrices device (Kent Scientifics) for amplicon sequencing and PCR.

For RNA hybridization, fresh mouse brain sections were stained using the RNAScope Multiplex V1 kit (ACDBio) with customized SaCas9-ATTO550 C2 and NeuN-Alexa Fluor488 C1, GFAP-ATTO647 C3 controls, following manufacturer’s instruction. Stained sections were visualized using Nikon N-STORM spinning-disk confocal microscope in the Confocal and Specialized Microscopy Shared Facility of the Herbert Irving Comprehensive Cancer Center at Columbia University.

For amplicon sequencing and PCR quantification, tissue was homogenized in 1× DNA/RNA Shield solution (Zymo Research) on ice followed by total RNA and DNA extraction using the Quick-DNA/RNA Miniprep kit (Zymo Research). AAV quantification in tissue was done by qPCR using the standards and primers from Takara’s quantification kit. To determine the gene editing performance, amplicon NGS was done followed by the same method used for *in vitro* validations. Results were analyzed using CRISPresso2, and the modified % obtained from the analysis was reported as gene editing rate %.

### Gene editing analysis in reporter mouse models

Adult female and male reporter mice (Ai9 or TLR2; 9-10 weeks old; JAX Lab) were given with 2×10^12^ vg/mouse AAV9 encoding SaCas9 and reporter-specific gRNAs (Supplementary Table 1) under the FUS array treatment. At week 3 post-administration, the mouse brain was harvested and processed for histological analysis by following the SCGE SATC standardized protocol. Briefly, under anesthesia, the mouse was transcardially perfused with cold 1×PBS followed by cold 4% paraformaldehyde (PFA)/PBS solution. After harvesting, the brain tissue was incubated in 4% PFA/PBS solution at 4°C with gentle shaking for 4-5h. After 2 washes with cold PBS, the tissue was incubated overnight in 30% sucrose/PBS at 4°C with gentle shaking. The sample was then embedded in OCT and serially sectioned with a thickness of 40 μm. Staining was done using the free-floating immunohistochemistry method. Sections were first washed with 1×TBS with 0.05% Triton X-100 (Sigma) and then incubated with 1×TBS containing 0.05% Triton X-100, 5% normal donkey serum and 2% BSA for 2h. After blocking, sections were stained with primary antibodies overnight. For TLR2 sections, sections were stained with both mouse anti-NeuN (1:1,000; Abcam) and rabbit anti-RFP antibodies (1:500; Thermo-Fisher), while only anti-NeuN antibody was used for Ai9 sections.

Following primary antibody staining, 3 washes with 1×TBS with 0.05% Triton X-100 were carried out, and 10 min for each wash. Sections were then incubated with the secondary antibodies at RT for 2h. Alexa Fluor 555 donkey anti-rabbit (1:1,000; Thermo-Fisher) and Alexa Fluor 488 donkey anti-mouse (1:1,000; Thermo-Fisher) were used for TLR2 sections, while only the donkey anti-mouse secondary antibody was used for Ai9 sections. After the incubation with the secondary antibody, three 10-min PBS washes were performed followed by counterstaining with DAPI (0.5 μg/mL in PBS; Thermo-Fisher) for 12 mins. Three additional PBS washes were done afterwards. Stained sections were then mounted onto the slides with Prolong Diamond antifade mounting media (Thermo-Fisher) and cued for 24 h at dark.

For visualization at Columbia, sections were scanned using Leica Aperio Versa8 multichannel fluorescence slide scanner under the 10× magnification in the Digital and Computational Pathology Laboratory at New York-Presbyterian Hospital for edited volume determination, and the hippocampus ROI images were scanned using Nikon N-STORM spinning-disk confocal microscope under the 20× magnification. At JAX, sections were scanned using Leica DMI8 microscope for whole-brain imaging, while Leica confocal microscope for hippocampus ROI imaging.

### Imaging quantification

For edited brain volume calculation, scanned whole brain images were analyzed using Image J (version 1.53t). Each brain image was divided into FUS and control parts. The total area of each part was obtained by applying the Triangle Threshold to generate binary images. The edited area (red) was identified from the binary image by applying the Yen or Moment Threshold. The edited ratio% was then determined by the ratio of edited area to total area. Furthermore, the edited volume was estimated by the integration of the red area in each slide. Briefly, the average red area in each slide was multiplied by the sum of the total thickness of slides and the total interval between slides:

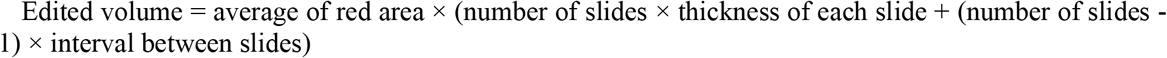

For edited neuron% in each ROI was quantified by QuPath (version 0.3.2). All the cells were first captured by the DAPI signal, and then two classifiers (green for neurons and red for TagRFP+ or TdTomato+ cells) were applied to obtain the cell number in each group. The ratio of the number of green- and red-colocalized cells to that of neurons (green) was reported as the edited neuron%.

### Experimental correction for the quantification in TLR2 mice

As the TagRFP in TLR2 mice could be only activated by the +1/-2 frameshifts, to better estimate the gene editing performance, we used the amplicon sequencing results obtained from the whole FUS-targeted brain hemisphere samples to calculate the correction factor for Figure 2h. The correction factor was defined as the ratio of total indels to +1/-2 indels (Extended Fig. 5b).

### Lighsheet imaging and quantification

For lightsheet imaging, we followed our previously published protocol^20^ to process the mouse brain tissues. After the PBS/PFA perfusion, brain tissue was first washed with PBS and then incubated with the hydrogel monomer solution (1% acrylamide, 0.0125% bisacrylamide, thermal initiator VA-044 in 4% PFA/PBS solution) overnight at 4°C. Afterwards, the sample was degassed, followed by hydrogel polymerization at 37°C for 4-5h with gentle shaking. Brain tissue was transferred to the clearing buffer containing 4% sodium dodecyl sulfate in 0.2 M boric buffer at pH 8.5 and incubated for ∼4 weeks. Once the tissue became clear, it was washed with 0.2M boric acid buffer (pH 8.5) with 0.1% Triton X-100. We then used our homemade CLARITY-optimized light-sheet microscopy^20^ for sample visualization. The sample was mounted using RapiClear (SunJin Lab) in a 10 mm light-path quartz cuvette (refractive index∼1.458; FireFlySci) and visualized using a 16×/0.4NA Objective (demagnified to 10×; ASI). The acquired lightsheet images therefore came with a resolution of 0.65 μm per pixel on both x- and y-directions, while 5 μm per pixel in the z-direction. The tile images were stitched using ImageJ, which generated roughly fifteen hundred 6,000×4,000 slides for each brain sample. To determine the edited volume, we first tested the readout of different thresholding methods. Among them, the Triangle algorithm was the best in capturing signals, but its triangle thresholding was not stable. To address this issue, we plotted the histogram of the threshold values and found the two values by fitting the histogram with two Gaussian kernels. This helped us to identify the thresholds corresponding to the autofluorescence and the real TdTomato signals (Supplementary Fig. 1). We therefore applied these two values to each slide for masking. Binary images could be then created, and the positive values were collected for quantification.

### Statistical Analysis

Unless specified, an unpaired, two-tailed Student t-test was used for the comparison of the data with only two groups, while one-way ANOVA with Tukey post-hoc analysis was used for multiple comparisons. Data are presented as mean ± standard error of the mean (SEM), and significance are presented as *, *p* < 0.05; **, *p* < 0.01; ***, *p* < 0.001; ****, *p* < 0.0001; and n.s., no significance.

## Acknowledgements

Brain structures in Figure 2b were drawn using BioRender. We would like to acknowledge the technical support from Mr. Zhong Wang of the Digital and Computational Pathology Laboratory at New York-Presbyterian Hospital, Dr. Theresa Swayne and her team of the Confocal and Specialized Microscopy Shared Resource at Columbia Herbert Irving Comprehensive Cancer Center, funded in part through the Center Grant P30CA013696. This work was supported by NIH SCGE Program (UG3NS115598, K.W.L., E.E.K., A.C; U42OD026635, S.A.M., C.M.L.) and the Career Development Award from American Society of Gene & Cell Therapy (ASGCT2021003, Y.-H.L.). The content is solely the responsibility of the authors and does not necessarily represent the official views of the funders.

## Extended Figures

**Extended Fig. 1.**
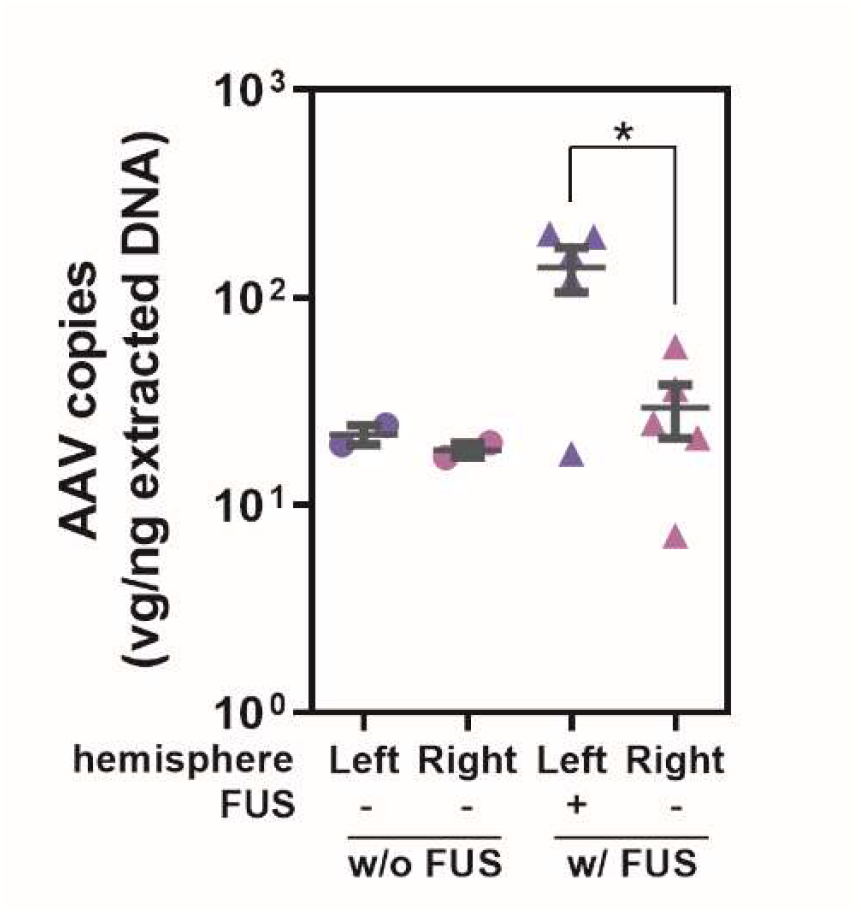
Spatiotemporal delivery of AAV/SaCas9 vector. Deposition of SaCas9/AAV9 vector in the different hemisphere at week 2 post-administration. Both FUS and control groups received an AAV dose of 2×10^11^ gc/mouse (*N*=5 for the FUS group and *N*=2 for the control group without FUS, adult male C57BL/6).

**Extended Fig. 2.**
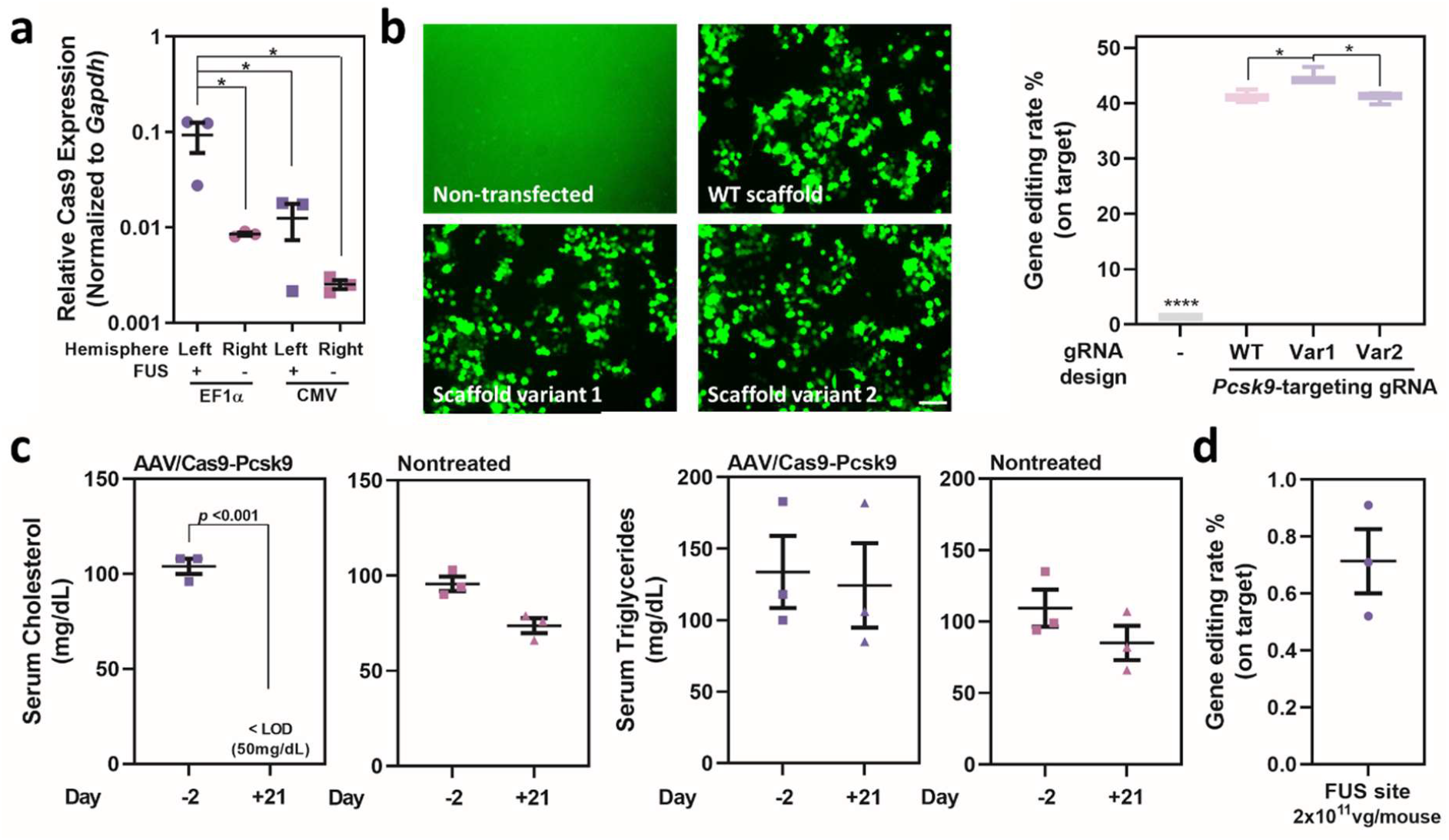
SaCas9 Vector optimization. **a**, SaCas9 expression level in the brain when mice given with EF1α- or CMV-driven SaCas9 AAV9 vector. Adult male C57BL/6 mouse was intravenously given the AAV9/SaCas9 vector in a dose of 2×10^11^ gc/mouse, and FUS-mediated BBB opening was performed in the left hemisphere. Mouse brain was dissected at week 3 post-administration for RNA extraction and downstream qPCR analysis (*N*=3 for each group). **b**, *In vitro* gRNA scaffold optimization. Mouse Neuro-2a cell was transfected with *Pcsk9*-targeting SaCas9 AAV9 vector encoding wild-type or the engineered variant and EGFP plasmid. At 48h post-transfection, genomic DNA was extracted, and the target *Pcsk9* locus was amplified for amplicon sequencing analysis. Amplicon sequencing results were analyzed using CRISPresso2, and the modified% was reported as the on-target gene editing rate% (two biological repeats were done for each group). Data are presented in a Box & Whiskers plot with minimum to maximum values **c**, Change of lipid profile after systemic administration of *Pcsk9*-targeting SaCas9 AAV vector with a dose of 2×10^11^ gc/mouse (*N*=3 for both FUS and nontreated groups). Analysis of the serum lipid panel was done by the Diagnosis Lab in the Institute of Comparative Medicine at Columbia University. **d**, *Pcsk9* gene editing rate at the FUS-targeted brain hemisphere when given with a dose of 2×10^11^ gc/mouse. The mouse brain was dissected at week 3 post-administration for genomic DNA extraction, and the target *Pcsk9* locus was amplified for amplicon sequencing analysis. Amplicon sequencing results were analyzed using CRISPresso2, and the modified% was reported as the on-target gene editing rate% (*N*=3).

**Extended Fig. 3.**
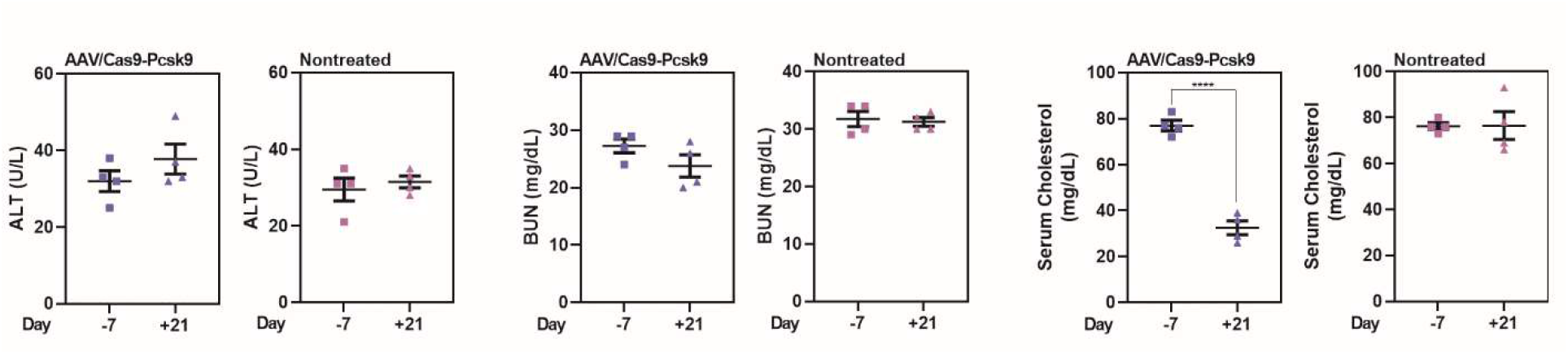
Change of serum biochemistry indicators when mice were given a higher SaCas9 AAV dose. Adult male C57BL/6 mice were intravenously given *Pcsk9*-targeting AAV9/SaCas9 (10^12^ gc/mouse), and FUS-mediated BBB opening was done in the left hemisphere. Serum was collected on day 7 prior to administration and week 3 post-administration (*N*=4 for both FUS and nontreated groups). Serum analysis was done by IDEXX.

**Extended Fig. 4.**
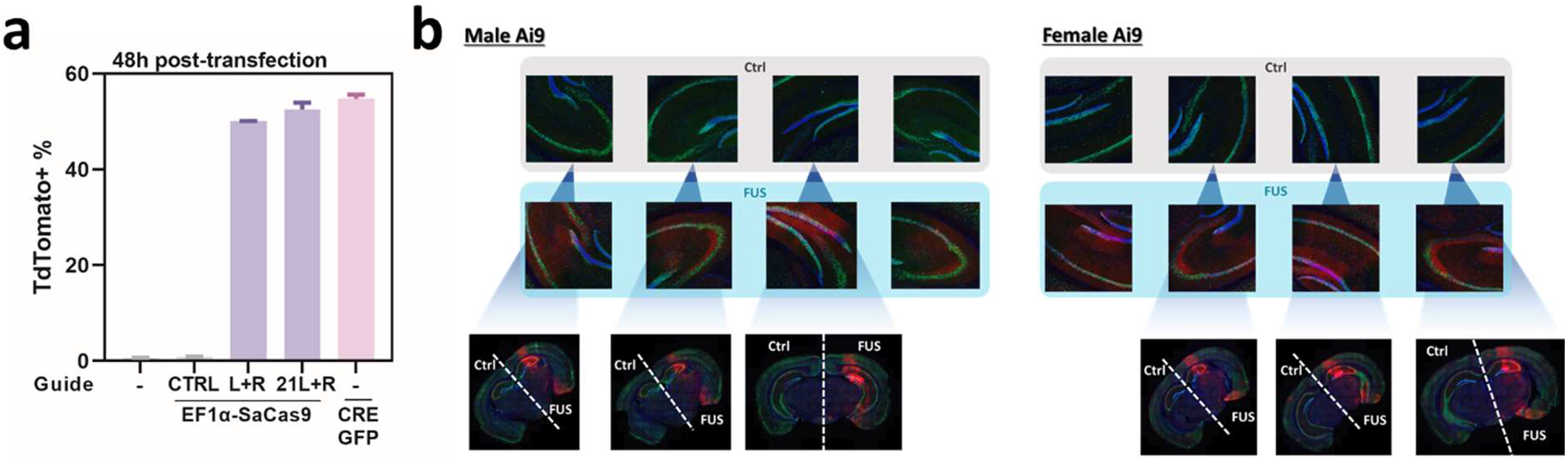
Validation in Ai9 reporter model. **a**, *In vitro* validation of the Ai9-targeting dual guide vector. HEK293T-Ai9 reporter cell was transfected with the dual guide SaCas9 AAV vector encoding either control (CTRL), Ai9-L+Ai9-R (L+R) or 21mer version of Ai9-L+Ai9-R (21L+R) gRNAs. At 48h post-transfection, all the cells were harvested and TdTomato+% of the whole cell population was determined by FACS (two biological repeats were done for each group). **b**, Confocal images used for edited neuron % determination in Figure 2e.

**Extended Fig. 5.**
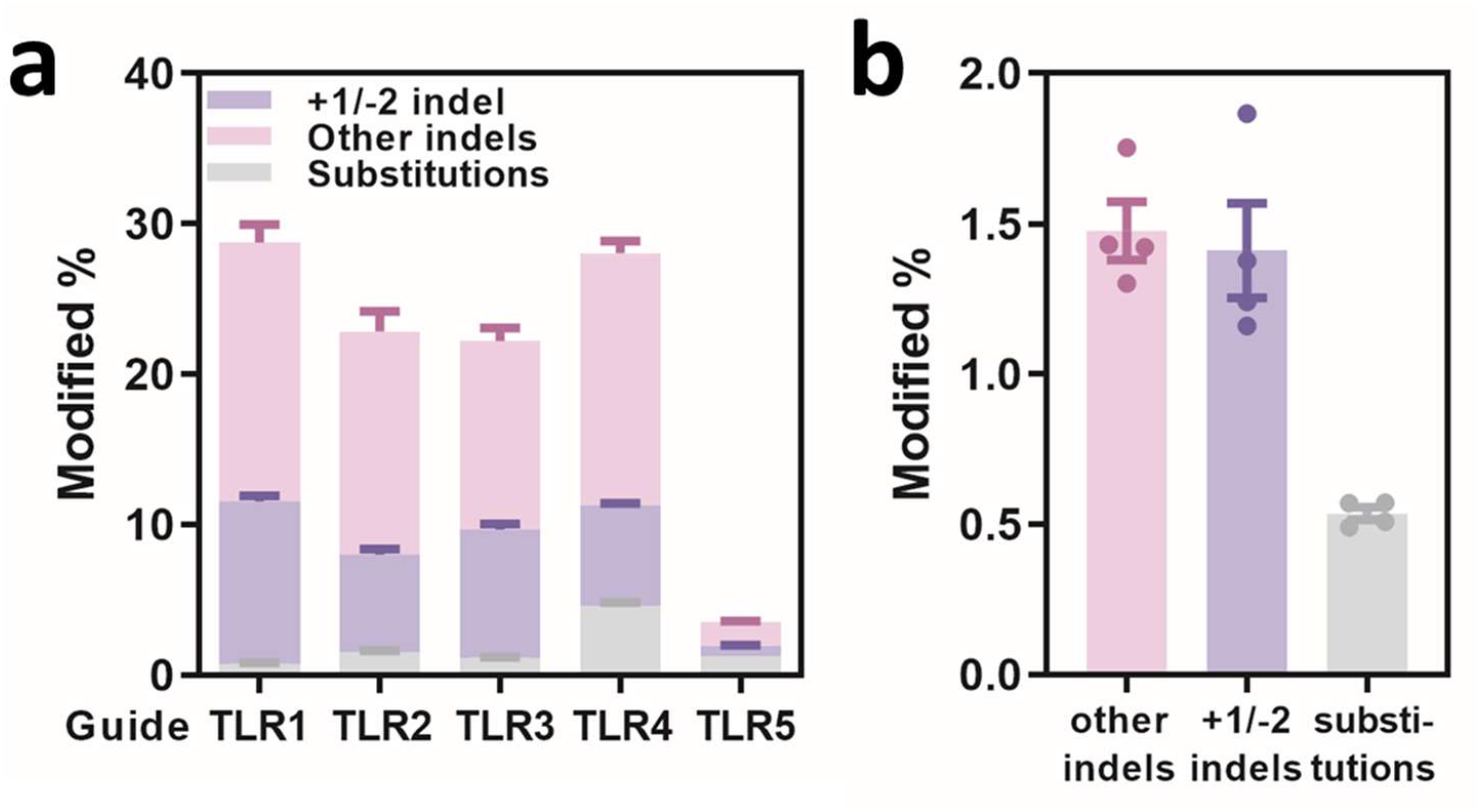
SaCas9 guide optimization and experimental correction for TLR2 reporter mouse model. **a**, TLR2 gRNA optimization. HEK293T-TLR2 reporter cell was transfected with SaCas9 AAV9 vector encoding each TLR2-targeting guide candidates. At 48h post-transfection, genomic DNA was extracted, and the target TLR2 locus was amplified for amplicon sequencing analysis. Amplicon sequencing results were analyzed using CRISPresso2 (two biological repeats were done for each group). **b**, Amplicon sequencing to determine the correction factor for gene editing performance analysis in TLR2 mice. Adult TLR2 mice were intravenously given the TLR2-targeting AAV9/SaCas9 vector in a dose of 2×10^12^ gc/mouse, and FUS-mediated BBB opening was performed using the array system at the left hemisphere. The whole targeted mouse hemisphere was dissected at week 3 post-administration, and the target TLR2 locus was retrieved by PCR followed by amplicon sequencing for analysis. Amplicon sequencing results were analyzed using CRISPresso2 (*N*=4 and male TLR2 mice used for this experiment).

