## Supplementary Materials for "Focused ultrasound-mediated brain genome editing"

**This document includes**

Supplementary Fig. 1. Threshold determination for light sheet image quantification.
Supplementary Table 1. The guides and gRNA backbones used in this study.
Supplementary Table 2. The primers used in this study.

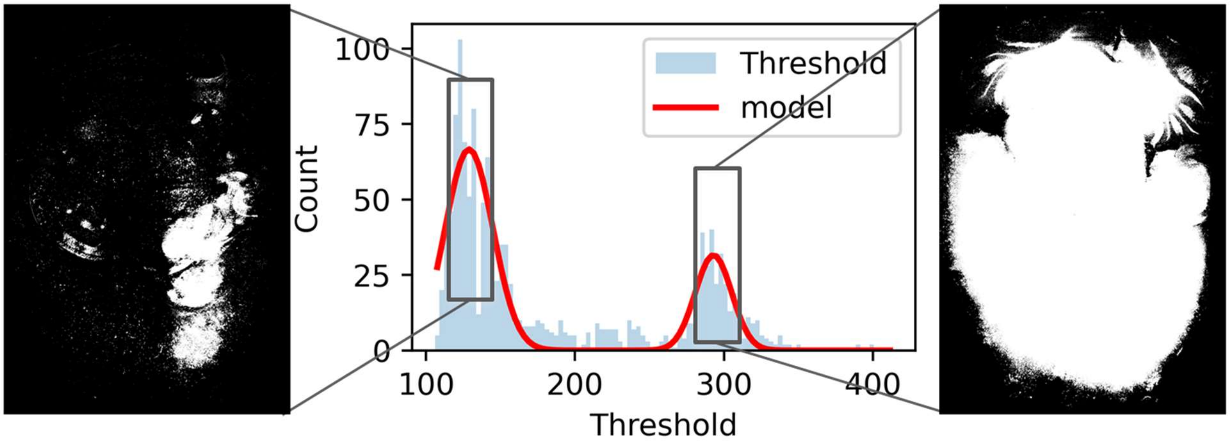


**Supplementary Fig. 1 | Threshold determination for light sheet image quantification.** A representative histogram showing the frequency distribution of signals from one set of the lightsheet image sections and the curve fitting to find the Gaussian kernels for determining the threshold for downstream quantification for Figure 2b.

**Supplementary Table. 1 | The guides and gRNA backbones used in this study.**

| ID# | target | sequence | used for | ref |
|---|---|---|---|---|
| Pcsk9 | *Pcsk9* | GCCGCTGACCACACCTGCCAG | Fig 1cde, Extended Fig 1,2,3 | 1 |
| Ai9-L | Ai9 | CTCTAGAGTCGCAGATCCTC | Extended Fig 4a | 2 |
| Ai9-R | | ACGAAGTTATATTAAGGGTT | | |
| Ai9-L21 | | CCTCTAGAGTCGCAGATCCTC | Fig. 2bcdef, Extended Fig 4 | |
| Ai9-R21 | | TACGAAGTTATATTAAGGGTT | | |
| TLR1 | TLR2 | ACGGCAACTACAAGACCCGCG | Fig 2gh, Extended Fig 5 | |
| TLR2 | | TACGTCGCCGTCCAGCTCGAC | Extended Fig 5a | |
| TLR3 | | GCCATGCCCGAAGGCTACGTC | | |
| TLR4 | | GAAGCACTGCAGGCCGTAGCC | | |
| TLR5 | | CACAAGTTCAGCGTGTCCGGC | | |
| wild-type gRNA backbone | | GTTTTAGTACTCTGGAAACAGAATCTACT AAAACAAGGCAAAATGCCGTGTTTATCT CGTCAACTTGTTGGCGAGA | Fig 1c. Extended Fig 1,2ab | 1 |
| gRNA backbone Var1 | | GTTATAGTACTCTGGAAACAGAATCTACT ATAACAAGGCAAAATGCCGTGTTTATCTC GTCAACTTGTTGGCGAGA | Fig 1de, Fig 2 Extended Fig 2,3,4,5 | 3 |
| gRNA backbone Var2 | | GTTTAAGTACTCTGTGCTGGAAACAGCAC AGAATCTACTTAAACAAGGCAAAATGCC GTGTTTATCTCGTCAACTTGTTGGCGAGA | Extended Fig 2ab | 2 |

**Supplementary Table. 2 | The primers used in this study.**

| primer | sequence | used for |
|---|---|---|
| *Pcsk9* amplicon seq | TATGAGGGTGCCGCTGACT | Fig 1f Extended Fig 2bd |
| | AGGGTCACCATCACCGACT | |
| TLR2 amplicon seq | AACCATGTTCATGCCTTCTTCTTT | Extended Fig 5 |
| | GATATAAGCCTGCCCAGAAGACT | |
